# Candidate Genes Associated with Parity Mode Evolution are Under Selection in Oviparous and Viviparous Snakes

**DOI:** 10.1101/2023.10.22.563505

**Authors:** X Maggs

**Affiliations:** American Museum of Natural History, Department of Herpetology; University of Missouri, Bond Life Sciences Center

**Keywords:** Reproductive Modes, Parity Modes, Snake Genomics, Oviparity, Viviparity, Diversifying Selection

## Abstract

Over 100 divergent lineages of squamates are viviparous (live-bearing) while others are oviparous (egg-laying). Estimated origins of oviparity and viviparity across the squamate phylogeny are a subject of scientific debate. Identifying genomic changes that influence parity mode evolution can provide insights on the adaptive nature of both parity modes. Using BUSTED in the HyPhy suite and whole genomes of 27 species of snakes, I test for gene-wide episodic diversifying selection (EDS) in 1,587 candidate genes. The genomes represent 14 viviparous and 13 oviparous species across four families (Viperidae, Elapidae, Natricidae, and Pythonidae), including eight new genomes from *Lachesis muta, Lachesis stenophrys, Agkistrodon contortrix, Trimeresurus albolabris, Gloydius halys, Ophryacus undulatus, Ovophis monticola*, and *Calloselasma rhodostoma*. Candidate genes were chosen based on evidence for their involvement of seven physiological processes expected to change during evolutionary transitions between parity modes. EDS was measured across candidate gene trees under different foreground branch tests: 1) all viviparous branches, 2) viviparous Viperidae branches, 3) viviparous Elapidae and Natricidae branches, 4) all oviparous branches, and 5) oviparous Viperidae branches (estimated reversals). Over 16% of tests found evidence for EDS, translating to 533 genes with EDS associated with parity mode. Some genes had significant EDS in multiple tests. However, the ‘all oviparous’ test identified a 2.8 to 5.9-fold increase in the number of unique genes with EDS compared to other foreground tests. These results suggest that EDS may influence the evolution of both oviparity and viviparity. Future research using population-level data from snakes may further elucidate the role of selection in parity mode evolution.

## Introduction

Viviparity (live birth) is estimated to have originated more frequently in squamates (snakes and lizards) than in all other vertebrate groups combined (Blackburn, 1992, 2015b). This makes squamates an opportune system for comparative research on the evolution of parity modes, oviparity (egg-laying) and viviparity. A wealth of data on uterine gene expression and functions during gestation and gravidity are now accessible thanks to decades of work—medical and veterinary research on pregnancy, evolutionary research on placental development in mammals, agricultural research on avian egg production, and comparative evolutionary genomics studies on closely related or reproductively bimodal lizards (Brandley et al., 2012; Cornetti et al., 2018; Foster et al., 2020, 2022; Gao et al., 2019; Griffith et al., 2016, 2017; Jonchere et al., 2010; Recknagel et al., 2021; Whittington et al., 2018; Xie et al., 2022).

Previous research has questioned whether independent viviparous lineages utilize the same molecular toolkit to support viviparity (Van Dyke et al., 2014). A genome wide association study in a reproductively bimodal lizard, *Zootoca vivipara*, revealed that eggshell thickness is genetically determined (Recknagel et al., 2021). This study identified 1,621 candidate genes with signatures of divergent selection in oviparous vs viviparous individuals (Recknagel et al., 2021). Over three hundred of these were differentially expressed in utero during gestation or gravidity (Recknagel et al., 2021), underscoring a link between gene expression and selection. Gao et al., 2019 also identified four genes (*C7, NKTR, NBEAL2, PTX*2) with amino acid substitutions in the same position across four species of viviparous skinks—suggestive of positive selection. Recent findings also reveal that shared parity/nutritional modes are not associated with convergent gene expression profiles in amniotes (Foster et al., 2022). This study concluded that the toolkits available to support parity mode evolution are restricted in a clade-specific fashion (Foster et al., 2022). Ancestral parity mode of a clade likely plays a role in this restriction.

Differing findings on the ancestral history of squamate parity modes suggest either viviparity evolved frequently or there was an early origin of viviparity that reversed back to oviparity many times (Blackburn, 1999, 2015a; Griffith et al., 2015; Harrington & Reeder, 2017; Pyron, 2015; Pyron & Burbrink, 2014, 2015). Discussion of ancestral parity modes of squamates is beyond the scope of this study, however, the ancestral state impacts the reproductive genomics/physiology of descendants. Parity mode evolution in snakes may be particularly unique given one study found a 90% probability of viviparity at the MRCA of all snakes (Harrington & Reeder, 2017). Another recent study found viviparous squamates had a significantly reduced number of intact avian eggshell-specific proteins compared to oviparous squamates, a pattern especially apparent in snakes (Xie et al., 2022). This makes snakes an interesting system for studying the evolution of parity modes. For this study, eight new genomes of pit vipers were sequenced— *Lachesis muta, Lachesis stenophrys, Agkistrodon contortrix, Trimeresurus albolabris, Gloydius halys, Ophryacus undulatus, Ovophis monticola*, and *Calloselasma rhodostoma* (Figure 1). This includes two species from the genus *Lachesis*, a genus phylogenetically supported as a reversal back to oviparity (Fenwick et al., 2012).

**Figure 1.**
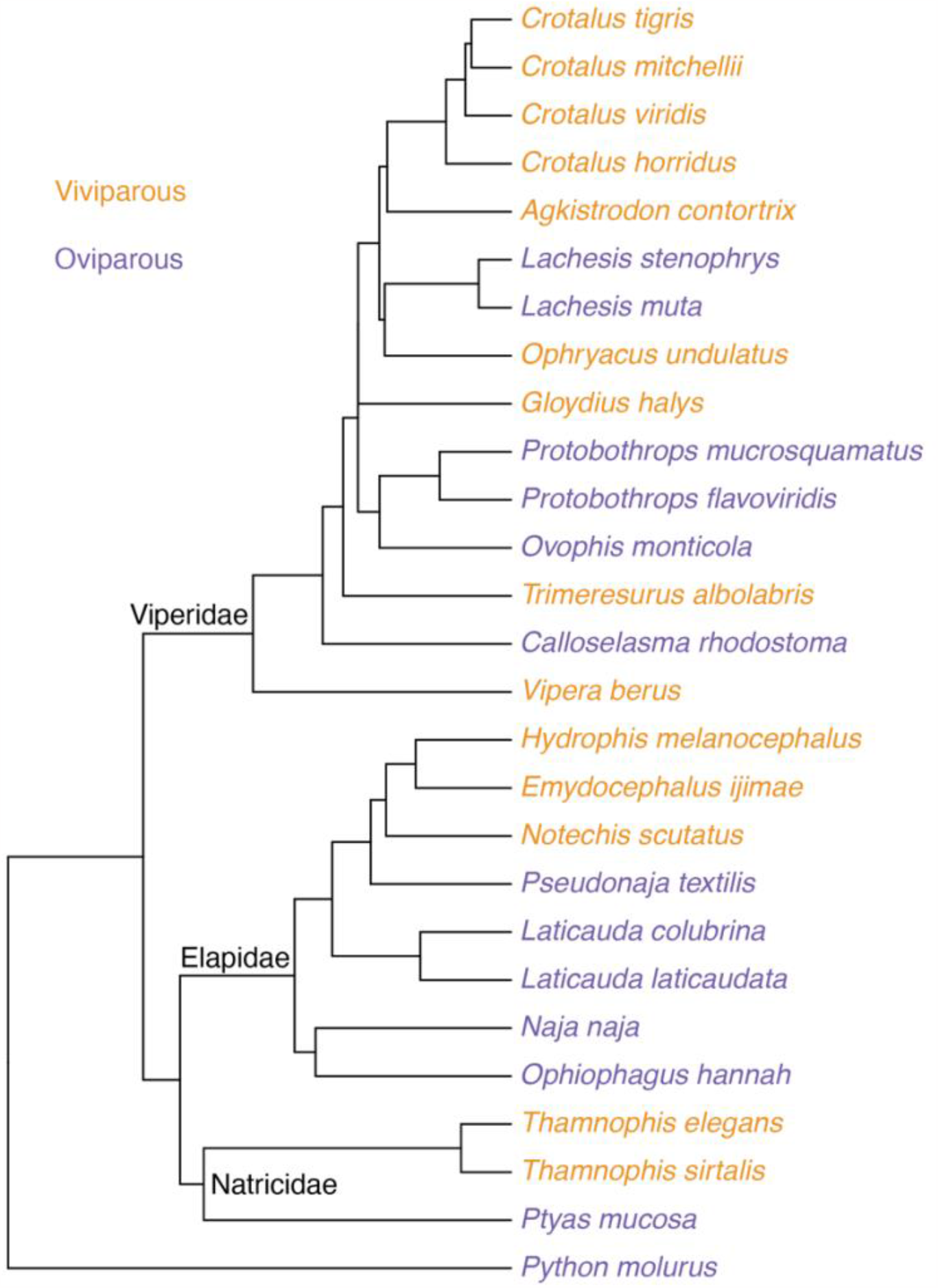
Phylogenetic relationships of taxa in study. Adapted from Pyron & Burbrink 2014.

Genes in a toolkit for viviparity should influence processes expected to changes to during transitions between parity modes—1) length of embryonic retention—only viviparous mothers retain the embryo for the entirety of development (Murphy & Thompson, 2011; Packard, Tracy, & Roth, 1977; Thompson & Speake, 2006); 2) eggshell deposition—viviparous embryos generally do not have an eggshell (Blackburn, 2015b; Heulin et al., 2005; Packard, Tracy, & Roth, 1977; van Dyke, Brandley, & Thompson, 2014b); 3) water provisioning; 4) gas provisioning; 5) nutrient provisioning; 6) development of a chorioallantoic placenta—viviparous embryos require, at minimum, a simple chorioallantoic placenta for water, gas and/or nutrient provisioning (Blackburn, 1992a, 2015b; Guillette, & Guillette, 1993; Thompson, Stewart, & Speake, 2000; Thompson & Speake, 2006b) maternal-fetal immune dynamics—viviparous gestation supports maternal tissue exposure to foreign fetal tissue and vice versa, which violates expectations of immune tolerance known from the peripheral body (Chavan et al., 2017; Graham et al., 2011; Hendrawan et al., 2017; Medawar, 1953). For this study, a large candidate gene list was developed to test for signatures of episodic diversifying selection (EDS), diversifying selection that occurs in a subset of branches.

A shared genetic toolkit for viviparity supported through EDS from an oviparous ancestor should meet two expectations—1) EDS in candidate genes should be relatively uncommon in oviparous branches, and 2) EDS of candidate genes in viviparous branches should be more enriched for the following functions—length of embryonic retention, water provisioning, gas provisioning, nutrient provisioning, development of a chorioallantoic placenta, and maternal-fetal immune dynamics—compared to candidates under selection in oviparous branches.

Using gene trees from candidate genes associated with parity mode evolution, I test whether there are higher likelihoods of EDS within 1) terminal viviparous foreground branches (hereafter all viviparous) and 2) terminal oviparous foreground branches (hereafter all oviparous), compared to the background. I also did this in a clade specific way, setting the foreground to 3) terminal viviparous viperid branches (hereafter viviparous viperids), 4) viviparous natricids and elapids (hereafter viviparous natricids and elapids), and 5) oviparous viperids (hereafter oviparous viperids). To test for EDS in any branch of the gene trees, I also ran a test with 6) no foreground branches set (hereafter all branches).

## Methods

### Taxonomic sampling and whole genome assembly

To increase the phylogenetic coverage of viperid snakes with whole genome data for this study, I obtained tissue or blood samples from the Texas Natural History Collection (TNHC) at the Texas Memorial Museum; the Department of Herpetology at the California Academy of Sciences (CAS); the Division of Herpetology at the University of Kansas (UK); the Dallas Zoo; and the Department of Herpetology at the American Museum of Natural History, stored in the Ambrose Monell Cryo Collection (AMCC) (Table 1). High molecular weight (HMW) DNA from tissues or blood was extracted using the MagAttract DNA kit. Six genomes were sequenced with the New York Genome Center using the Illumina Truseq library prep and Hiseq X sequencing platform (Table 1). For the Hiseq X genomes, reads were quality checked with FastQC version 0.11.5, trimmed with Trimmomatic version 0.39. Two more genomes for *Lachesis muta* and *Agkistrodon contortrix* were sequenced using 10X Chromium linked reads and assembled as a first pass with Supernova version 2.0. Nineteen other published genomes were used. Ten of these are chromosome-level genome assemblies, used without any changes (Table 1).

**Table 1.** Whole Genome Dataset, Tissue Loans and Published Assemblies

| Taxa | Genome Accession # | Tissue ID | Sequencing Method<sup>A</sup> | Genome Alignment Reference |
| --- | --- | --- | --- | --- |
| *Crotalus tigris*<sup>B</sup> | GCA_016545835.1 |  |  |  |
| *Crotalus mitchellii* | GCA_000737285.1 |  |  | *Crotalus viridis* |
| *Crotalus viridis*<sup>B</sup> | GCA_003400415.2 |  |  |  |
| *Crotalus horridus* | GCA_001625485.1 |  |  | *Crotalus viridis* |
| *Agkistrodon contortrix* |  | AMNH: FTB3978 | 10x chromium linked reads | *Crotalus viridis* |
| *Lachesis stenophrys* |  | TNHC: UTA-R 25598 | Truseq & Illumina HiSeqX | *Crotalus viridis* |
| *Lachesis muta* |  | Dallas Zoo female, blood draw | 10x chromium linked reads | *Crotalus viridis* |
| *Ophryacus undulatus* |  | AMCC: 118187 | Truseq & Illumina HiSeqX | *Crotalus viridis* |
| *Gloydius halys* |  | UK: 331042 | Truseq & Illumina HiSeqX | *Crotalus viridis* |
| *Protobothrops mucrosquamatus*<sup>B</sup> | GCA_001527695.3 |  |  |  |
| *Protobothrops flavoviridis*<sup>B</sup> | GCA_003402635.1 |  |  |  |
| *Ovophis monticola* |  | CAS: 224424 | Truseq & Illumina HiSeqX | *Crotalus viridis* |
| *Trimeresurus albolabris* |  | UK: 312226 | Truseq & Illumina HiSeqX | *Crotalus viridis* |
| *Calloselasma rhodostoma* |  | UK: 328570 | Truseq & Illumina HiSeqX | *Crotalus viridis* |
| *Vipera berus* | GCA_000800605.1 |  |  | *Crotalus viridis* |
| *Hydrophis melanocephalus* | GCA_004320005.1 |  |  | *Naja naja* |
| *Emydocephalus ijimae* | GCA_004319985.1 |  |  | *Naja naja* |
| *Notechis scutatus*<sup>B</sup> | GCA_900608555.1 |  |  |  |

| Taxa | Genome Accession # | Tissue ID | Sequencing Method<sup>A</sup> | Genome Alignment Reference: Accession # |
| --- | --- | --- | --- | --- |
| *Pseudonaja textilis*<sup>B</sup> | GCA_900608585.1 |  |  |  |
| *Laticauda colubrina*<sup>B</sup> | GCA_015471245.1 |  |  |  |
| *Laticauda laticauda* | GCA_004320025.1 |  |  | *Naja naja* |
| *Naja naja*<sup>B</sup> | GCA_009733165.1 |  |  |  |
| *Ophiophagus hannah* | GCA_000516915.1 |  |  | *Naja naja* |
| *Thamnophis elegans*<sup>B</sup> | GCA_009769535.1 |  |  |  |
| *Thamnophis sirtalis* | GCA_001077635.2 |  |  | *Thamnophis elegans* |
| *Ptyas mucosa*<sup>B</sup> | GCA_012654045.1 |  |  |  |
| *Python molurus bivittatus* | GCA_000186305.2 |  |  | *Crotalus viridis* |

Note:<sup>A</sup> Sequencing method only reported for species newly sequenced for this study. <sup>B</sup> Denotes species with published genome assemblies that were used as published.

**Table 2.** Number of Candidate Genes^*^ per Functional Cluster

| Cluster | Subcluster | # Gene IDs |
| --- | --- | --- |
| Reproductive Physiology | Eggshell formation | 373 |
|  | Embryonic retention | 304 |
|  | Placentation | 644 |
|  | Maternal-fetal immune dynamics | 303 |
|  | Embryonic oxygen demands | 148 |
|  | Embryonic calcium & nutrient provisioning | 51 |
|  | Water provisioning | 17 |
| Exploratory | Associated with viviparity by Cornetti et al. (2018) | 19 |
|  | DEGs in uterus during gestation in viviparous mammals | 28 |
|  | DEGs in uterus during gestation in viviparous squamates | 24 |
|  | Anole egg protein genes | 112 |
|  | Autoimmune disease association | 26 |

\*Several genes fall into more than one cluster.

To improve access to candidate gene sequences, I aligned nine scaffold-level published assemblies to a chromosome-level references. For these and the eight newly sequenced viperid genomes (Table 1), I used an alignment-based approach to build assemblies with the highest protein-coding content as possible. I used the closest related and highest quality genome assembly available at the time (2020) as a reference for read/scaffold mapping (Table 1). I aligned reads or scaffolds using bwa-mem in BWA version 0.7.15 with default settings. For newly sequenced genomes, fastq files for read one and read two were input input. BWA-mem generates a sam file as output with the alignment information. I sorted and identified mate coordinates using samtools fixmate function and samtools index to the bam file output in Samtools version 1.9.0. Variants and indels were called using the bcftools mpileup and bcftools call functions in bcftools version 1.13 at a maximum depth of 80. Indels were then normalized with bcftools norm with default parameters. Clusters of indels separated by 5 or fewer bases were filtered with bcftools filter. This file was then indexed with the tabix command in Samtools. Sites with zero coverage in the mapped reads, contigs or scaffolds were extracted using the genomecov command in bedtools version 2.26.0. A final consensus genome sequence was generated using bcftools consensus with the zero-coverage file and the vcf file containing single nucleotide polymorphisms and indels compared to the reference. Sequences comprised entirely with Ns were removed with sequence_cleaner.py (https://biopython.org/wiki/Sequence_Cleaner).

The final dataset includes 27 snake genomes, representing 14 viviparous and 13 oviparous species from the families Elapidae (n=8), Viperidae (n=15), Natricidae (n=3), and Pythonidae (n=1) as the outgroup (Figure 1). Genome assembly quality was assessed with Quast v5.0.2 (Table 3). BUSCO v 5.2.2 was used to assess the number of near-universal single copy orthologs present in the genomes, with the Tetrapoda database 10 and Metaeuk gene predictor (Manni et al., 2021) (Table 4).

**Table 3.** Summary Statistics of 27 Snake Genome Assemblies

| Taxa | Scaffold N50 | Total length | GC % | # Ns per 100 kbp |
| --- | --- | --- | --- | --- |
| *Crotalus tigris* | 2,106,662 | 1,611,396,967 | 39.88 | 0 |
| *Crotalus mitchellii*<sup>C</sup> | 179,897,795<br>(5,299) | 1,331,920,201 | 38.11 | 21,795 |
| *Crotalus viridis* | 179,897,795 | 1,331,924,206 | 38.86 | 6,199 |
| *Crotalus horridus*<sup>C</sup> | 180,762,042<br>(23,829) | 1,338,211,176 | 38.58 | 13,126 |
| *Agkistrodon contortrix*<sup>N</sup> | 181,255,822 | 1,341,670,516 | 38.56 | 16,805 |
| *Lachesis stenophrys*<sup>N</sup> | 180,658,615 | 1,337,374,174 | 38.98 | 9,958 |
| *Lachesis muta*<sup>N</sup> | 181,290,216 | 1,341,824,342 | 38.61 | 17,000 |
| *Ophryacus undulatus*<sup>N</sup> | 180,652,035 | 1,337,239,219 | 39.02 | 10,438 |
| *Gloydus halys*<sup>N</sup> | 180,731,309 | 1,337,818,961 | 38.91 | 12,841 |
| *Protobothrops mucrosquamatus* | 424,052 | 1,621,677,801 | 39.88 | 8,115 |
| *Protobothrops flavoviridis* | 467,050 | 1,380,220,834 | 39.41 | 3,282 |
| *Ovophis monticola*<sup>N</sup> | 180,783,314 | 1,338,151,417 | 38.99 | 12,442 |
| *Trimeresurus albolabris*<sup>N</sup> | 180,678,254 | 1,337,361,094 | 38.69 | 14,380 |
| *Calloselasma rhodostoma*<sup>N</sup> | 180,375,891 | 1,335,130,470 | 38.23 | 18,469 |
| *Vipera berus*<sup>C</sup> | 180,756,280<br>(126,452) | 1,337,313,601 | 38.12 | 32,420 |
| *Hydrophis melanocephalus*<sup>C</sup> | 224,770,497<br>(59,810) | 1,772,964,496 | 38.16 | 41,376 |
| *Emydocephalus ijimae*<sup>C</sup> | 224,840,317<br>(18,937) | 1,773,465,297 | 38.56 | 38,835 |
| *Notechis scutatus* | 8,497,191 | 1,619,909,044 | 39.62 | 4,957 |
| *Pseudonaja textilis* | 17,062,011 | 1,566,804,039 | 39.77 | 2,498 |
| *Laticauda colubrina* | 39,959,644 | 2,024,955,717 | 41.24 | 8,931 |
| *Laticauda laticauda*<sup>C</sup> | 224,731,386<br>(39,330) | 1,772,834,246 | 38.24 | 39,047 |
| *Naja naja* | 224,088,900 | 1,768,461,657 | 40.46 | 6,202 |
| *Ophiophagus hannah*<sup>C</sup> | 225,072,945<br>(241,519) | 1,775,496,769 | 38.96 | 33,723 |
| *Thamnophis elegans* | 100,851,885 | 1,672,189,689 | 40.95 | 2,284 |
| *Thamnophis sirtalis*<sup>C</sup> | 101,200,466<br>(647,592) | 1,677,804,592 | 39.70 | 37,366 |
| *Ptyas mucosa* | 15,963,960 | 1,707,539,727 | 40.55 | 3,344 |
| *Python molurus bivittatus*<sup>C</sup> | 179,913,841<br>(213,970) | 1,332,019,523 | 37.83 | 80,968 |

Note: <sup>C</sup> denotes previously published genomes aligned to chromosome-level reference. For these, numbers in parenthesis indicates value prior to alignment to genome reference. <sup>N</sup> denotes new genome assemblies generated in this study.

**Table 4.** BUSCO Scores across 27 Snake Genomes

| Taxa | % Complete | % Single | % Duplicates | % Fragments | % Missing |
| --- | --- | --- | --- | --- | --- |
| *Crotalus tigris* | 95.60% | 94.10% | 1.50% | 1.00% | 3.40% |
| *Crotalus mitchellii*<sup>C</sup> | 85.90% | 85.60% | 0.30% | 5.30% | 8.80% |
| *Crotalus viridis* | 86.70% | 86.30% | 0.40% | 5.00% | 8.30% |
| *Crotalus horridus*<sup>C</sup> | 85.90% | 85.40% | 0.50% | 5.50% | 8.60% |
| *Agkistrodon contortrix*<sup>N</sup> | 84.80% | 84.30% | 0.50% | 5.80% | 9.40% |
| *Lachesis stenophrys*<sup>N</sup> | 86.30% | 85.70% | 0.60% | 5.30% | 8.40% |
| *Lachesis muta*<sup>N</sup> | 84.90% | 84.40% | 0.50% | 5.70% | 9.40% |
| *Ophryacus undulatus*<sup>N</sup> | 86.50% | 86.10% | 0.40% | 5.10% | 8.40% |
| *Gloydus halys*<sup>N</sup> | 86.60% | 86.10% | 0.50% | 5.00% | 8.40% |
| *Protobothrops mucrosquamatus* | 92.40% | 90.90% | 1.50% | 3.70% | 3.90% |
| *Protobothrops flavoviridis* | 90.20% | 89.40% | 0.80% | 4.50% | 5.30% |
| *Ovophis monticola*<sup>N</sup> | 86.70% | 86.20% | 0.50% | 4.90% | 8.40% |
| *Trimeresurus albolabris*<sup>N</sup> | 86.80% | 86.40% | 0.40% | 5.00% | 8.20% |
| *Calloselasma rhodostoma*<sup>N</sup> | 86.40% | 86.00% | 0.40% | 5.10% | 8.50% |
| *Vipera berus*<sup>C</sup> | 82.70% | 82.20% | 0.50% | 6.50% | 10.80% |
| *Hydrophis melanocephalus*<sup>C</sup> | 88.90% | 88.30% | 0.60% | 3.70% | 7.40% |
| *Emydocephalus ijimae*<sup>C</sup> | 91.30% | 90.70% | 0.60% | 2.50% | 6.20% |
| *Notechis scutatus* | 89.60% | 87.70% | 1.90% | 3.90% | 6.50% |
| *Pseudonaja textilis* | 93.40% | 91.10% | 2.30% | 2.20% | 4.40% |
| *Laticauda colubrina* | 92.70% | 92.10% | 0.60% | 2.50% | 4.80% |
| *Laticauda laticauda*<sup>C</sup> | 91.50% | 90.90% | 0.60% | 2.70% | 5.80% |
| *Naja naja* | 91.90% | 90.90% | 1.00% | 2.50% | 5.60% |
| *Ophiophagus hannah*<sup>C</sup> | 90.60% | 89.90% | 0.70% | 3.10% | 6.30% |
| *Thamnophis elegans* | 92.10% | 90.60% | 1.50% | 2.20% | 5.70% |
| *Thamnophis sirtalis*<sup>C</sup> | 73.70% | 73.20% | 0.50% | 8.40% | 17.90% |
| *Ptyas mucosa* | 91.20% | 90.00% | 1.20% | 3.30% | 5.50% |
| *Python molurus bivittatus*<sup>C</sup> | 73.00% | 72.60% | 0.40% | 9.40% | 17.60% |

Note: <sup>C</sup> denotes previously published genomes aligned to chromosome-level reference. <sup>N</sup> denotes new genome assemblies generated in this study. Values represent scaffolds/contigs including gaps.

### Selecting candidate genes

The candidate gene list was established between 2017-2021 based on the literature at the time. The criteria for choosing candidates, the full list of candidate genes (n=1587), and their membership in the reproductive physiology clusters—eggshell formation, embryonic retention, placentation, maternal-fetal immune dynamics, embryonic gas demands, embryonic nutrient provisioning, or water provisioning—can be found in the supplementary material. Candidate genes were compiled largely from studies on the uterine gene expression profiles of oviparous and viviparous lizards during gestation and gravidity (Brandley et al., 2012; Foster et al., 2020; Gao et al., 2019; Griffith et al., 2016, 2017; Recknagel et al., 2021). Some candidate genes were not previously directly associated with one of the physiological clusters. Several studies on squamate uterine gene expression have associated different reproductive physiology clusters with enrichment of specific GO terms. When a gene was not specifically implicated in a reproductive physiology cluster, I categorized it into a cluster if it 1) is differentially expressed during gestation or gravidity in avian or squamate female reproductive tissue, and 2) enriches functional terms previously associated with the cluster. Many genes were associated with multiple reproductive physiology clusters. Justification for the categorization of each gene into a cluster, and associated citations are provided in the supplement.

In addition to the genes in the reproductive physiology clusters, I also tested for EDS in genes that are expressed, and have evolved rapidly, in the *Anolis carolinensis* egg (Alföldi et al., 2011), additional genes associated with the transition to viviparity (Cornetti et al., 2018) and genes expressed in utero during gestation in four or more species of squamates or mammals (Recknagel et al., 2022). I also included a small (n=26) exploratory set of genes associated with human autoimmune diseases (Zhernakova et al., 2009).

### Obtaining query sequences for candidate loci

To identify homologous candidate loci in the genomes, I first downloaded query sequences for 1871 candidate genes (supplementary material). I downloaded genome annotations for *Thamnophis elegans* (NCBI: GCF_009769535.1_rThaEle1.pri_genomic.gff*), Crotalus viridis (*Figshare: *CroVir_rnd1*.*all*.*maker*.*final*.*homologIDs_rename*.*gff), Notechis scutatus* (NCBI: Notscu_GCF_900518725.1_TS10Xv2-PRI_genomic.gff), and *Python molurus bivittatus* (NCBI: GCF_000186305.1_Python_molurus_bivittatus-5.0.2_genomic.gff) and pulled all CDS sequences of candidates from them using the gffread tool. For candidate genes that were not annotated in these genomes, I pulled them from the *Anolis carolinensis* (NCBI: GCF_000090745.1_AnoCar2.0_genomic.gff), then *Gallus* (NCBI: GCF_016699485.2_bGalGal1.mat.broiler.GRCg7b_genomic.gff), and finally from *Homo sapiens* (NCBI: Homsap_GCF_000001405.39_GRCh38.p13_genomic.gff) genomes. For genes with multiple isoforms, I used the longest sequence as the query.

### Downloading queries and aligning candidate genes

I used query sequences as input for Splign version 2.1.0 which identified homologous regions in the 27 genomes with e-value=0.01. Splign is a utility for computing spliced sequence alignments (Kapustin et al., 2008). The candidates recovered from the 27 genomes, were aligned to one another using MASCE, a multiple sequence alignment algorithm that accounts for frame shifts and stop codons (Ranwez et al., 2011). Based on the limitations of BUSTED input, I converted frameshift insertions (!) to gaps (-), removed stop codons from the ends of sequences, and removed sequences with internal stop codons prior to downstream analyses. Alignments with less than 10 sequences were removed from downstream analyses.

*Building gene trees*

Best-fit codon models were estimated with ModelFinder (Kalyaanamoorthy et al., 2017) from the Bayesian Information Criteria (BIC) score in IQ-TREE version 1.6.12 (Nguyen et al., 2015) using ultrafast bootstrap maximum likelihood estimation over 1,000 replicates (Thi Hoang et al., 2017). Gene trees were generated for the best-fit model for each gene and used as input for selection analyses in HyPhy version 2.5 (Kosakovsky Pond, et al., 2020).

### Episodic selection

To test specific hypotheses regarding foreground and background branches I tested for gene-wide EDS with BUSTED (Murrell et al., 2015). Gene trees with less than ten terminal branches were excluded from the dataset. BUSTED uses the Dn/Ds ratio across proportions of foreground and background branch-sites to identify gene-wide selection. A likelihood ratio test is run comparing a model with unrestricted Dn/Ds in foreground branches to a model restricted to Dn/Ds < 1. As this research was highly exploratory, and more phylogenetic sampling is needed to find conclusive results, tests with p < 0.05 for Dn/Ds > 1 are reported as significant. Precise p-values can be found in the supplementary material for further hypothesis testing. Significant results mean that there is evidence for EDS in at least one branch in the foreground.

I first ran BUSTED across the gene trees with no foreground specified (all branch test). Then to infer if EDS was occurring in association with parity mode evolution, I ran several foreground branch tests. To ascertain whether any genes are under diversifying selection in viviparous lineages specifically, I set foreground branches to terminal branches of gene sequences from viviparous taxa (all viviparous test). To determine if any genes have EDS within specific viviparous clades, I ran two tests, one setting foreground branches to all terminal branches representing viviparous viperid taxa (viviparous viperids test) and another set to terminal branches of viviparous natricids and elapids (viviparous natricids and elapids test). To determine if diversifying selection plays a role in maintaining or evolving oviparous reproduction, I ran a hypothesis test setting foreground branches to all oviparous terminal branches (all oviparous test). To test for EDS in oviparous viperid lineages, I set foreground to terminal branches of oviparous viperids (estimated reversals test). Internal branches were not used as foreground in any of these tests, other than the ‘all branch test’, because the state at internal branches is essentially unknown.

## Results

### Genome quality assessment

Statistics assessing genome quality and content are in Tables 4 and 5. All genomes had a scaffold N50 > two million base pairs except for *Protobothrops mucrosquamatus* and *Protobothrops flavoviridis* (Table 3). The GC content was similar, ∼40%, across the genomes. The genomes varied in terms of the number of Ns per 100 kbp. Draft genomes that were assembled through reference alignment had more Ns than some of the de novo assemblies, but the highest number of Ns per 100 kbp was found in Elapidae, Pythonidae and Natricidae genomes (Table 3). Regarding BUSCO scores, all genomes had over 82% complete BUSCOs except *Thamnophis sirtalis* and *Python molurus* which both had ∼73% (Table 4). Generally, percent duplicates were under 1%, but this was variable (Table 4). The highest percent of duplicates was in the *Pseudonaja textilis* genome, 2.3% (Table 4). All newly sequenced genomes had BUSCO % complete above 84%, in line with the 86.6% in their reference assembly, *Crotalus viridis*.

### Episodic diversifying selection

Cumulatively, 16.6% of BUSTED tests (n=1,577) returned significant results, of which 37% (n=586) have p-values less than 5.0e-05. This translates to 785 genes with significant signatures of diversifying selection, 533 of which are significant in the five foreground branch tests. The genes with signatures of EDS are distributed across reproductive physiology clusters (Figure 2). Several genes were significant in multiple tests (Figure 3).

**Figure 2.**
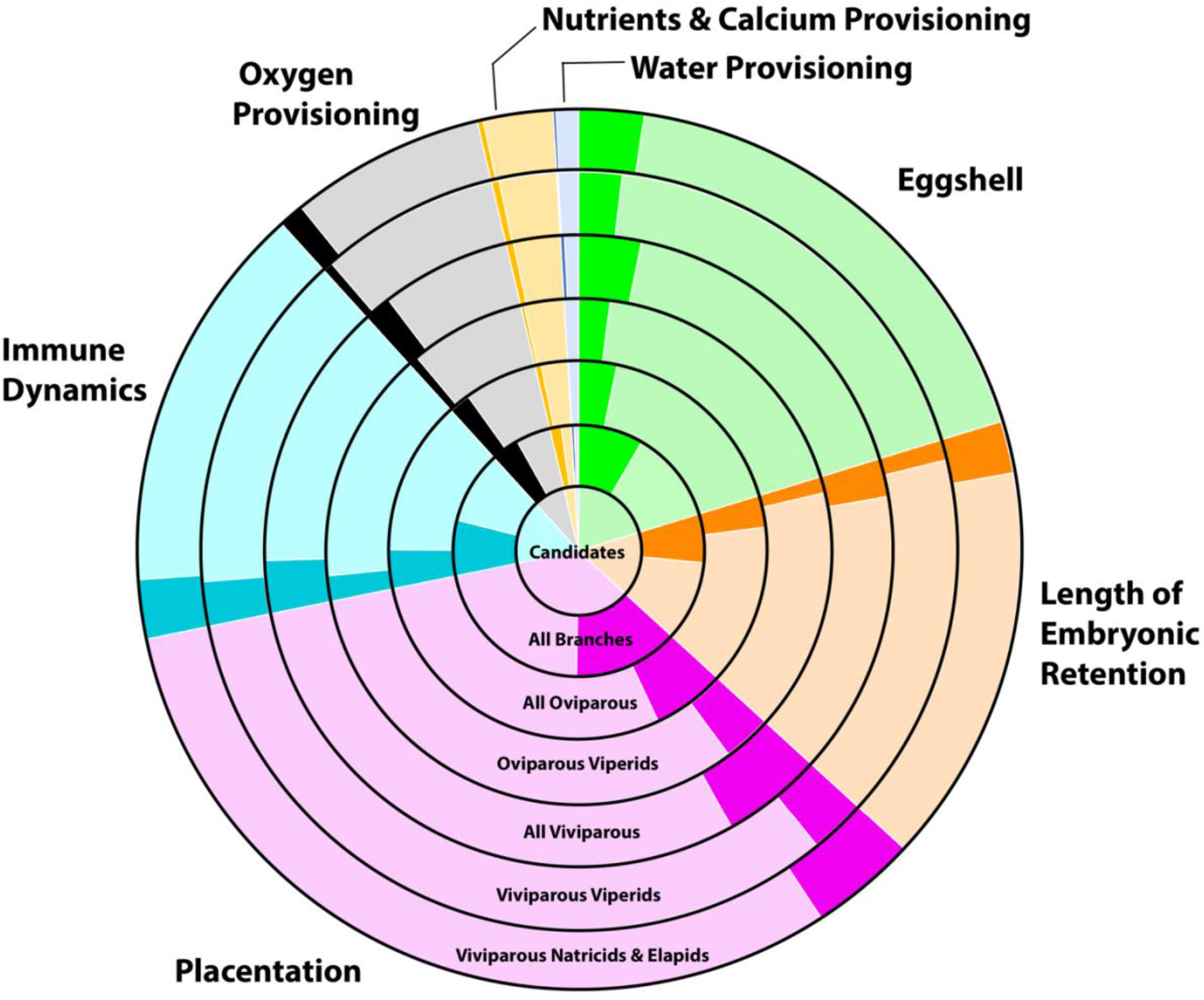
Proportion of genes with evidence for EDS across functional clusters for each foreground branch test.

**Figure 3.**
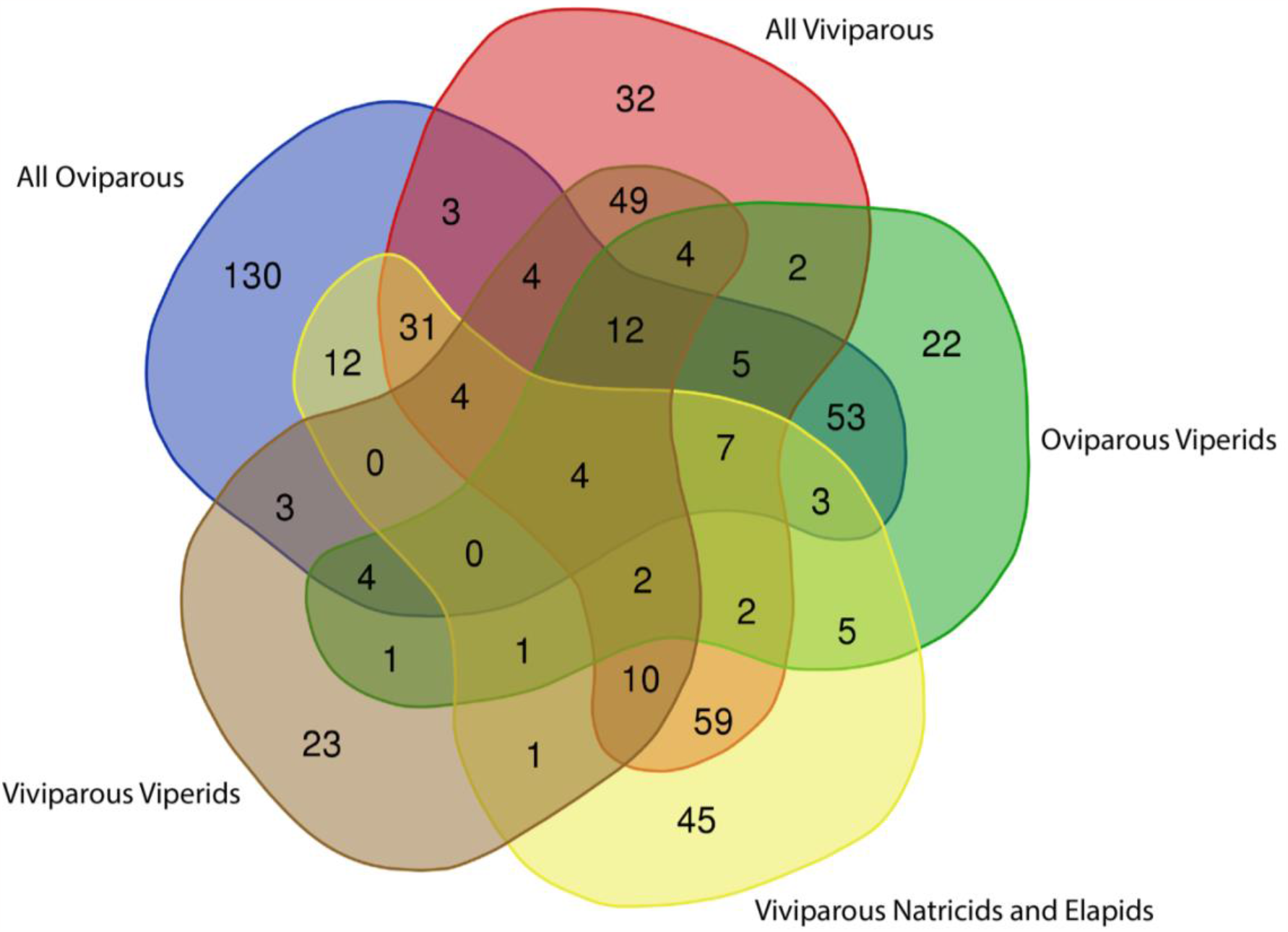
Overlaps of genes under EDS in five foreground branch tests.

Each test identified 100 or more genes with signatures of episodic diversifying selection. Over 40% of candidate genes (n=637) showed significant signatures of EDS in the all branch test (p < 0.05). In the all oviparous test, 275 genes showed significant signatures of EDS (p < 0.05). In the oviparous viperid test, 127 genes showed significant signatures of EDS (p < 0.05). In the all viviparous test, 230 genes had significant EDS (p < 0.05). In the viviparous viperids test, 122 genes had signatures of EDS (p < 0.05). In the viviparous natricids and elapids test, 186 genes showed significant signatures of EDS (p < 0.05).

### Comparing BUSTED tests

Subsets of genes were uniquely significant in specific foreground branch tests compared to the other four foreground branch tests (Figure 3)—all oviparous (n=130), all viviparous (n=32), oviparous viperids (n=22), viviparous natricids and elapids (n=45), and viviparous viperids (n=23). Pairwise overlaps were observed across tests, especially between all oviparous and oviparous viperids (n=53), all viviparous and viviparous natricids and elapids (n=59), and viviparous viperids and all viviparous (n=49).

## Discussion

Diversifying positive selection should not play a major role in the maintenance of an ancestral trait. Therefore, it is unexpected that a 2.8 to 5.9-fold increase was observed in the number of genes under EDS unique to the all oviparous test (n=130) compared to genes unique to viviparous foreground tests (Figure 2). Although further research is necessary to determine why these genes are under selection, the result highlights a putative relationship between diversifying selective and oviparous reproduction in snakes. This pressure in oviparous clades may reflect 1) the evolution of oviparity from viviparity, 2) the evolution of specifications to better support oviparous reproduction, or 3) the evolution of other processes unrelated to parity mode.

The five foreground branch tests identified over 530 genes with EDS. This suggests that both parity modes may be supported by EDS. However, there are no apparent relationships between viviparity and enrichment of genes with EDS in the following functional clusters—length of embryonic retention, water provisioning, gas provisioning, nutrient provisioning, development of the chorioallantoic placenta, and maternal-fetal immune response (Figure 2). At first glance, this suggests that a shared genetic toolkit for viviparity is not supported through EDS from an oviparous ancestor. However, given that selection plays an important role in both the evolution of constructive and regressive traits (i.e., gains and losses, respectively; Moran et al., 2023), broad scale patterns about the number and function of genes under selection in oviparous and viviparous lineages may not be inherently informative. For example, recent research on *Astyanax mexicanus* identified significant selective pressure on loci associated with regressive traits in multiple populations (e.g., eye loss, loss of pigmentation; Moran et al., 2023). Thus, genes that influence maternal-fetal immune dynamics, for example, may be under selection to support 1) enhanced immune-dynamics for viviparity, and 2) repressed immune dynamics for oviparity. The adaptive nature of oviparity is not well established, but some suggest either parity mode can be evolutionarily advantageous in different scenarios (Lee & Doughty, 1997; Packard, Tracy, & Roth, 1977; Rothchild, 2003; Shine & Bull, 1979a; Tinkle & Gibbons, 1977a).

Nonetheless, gene specific results are illuminating. A recent study linked the expression of *HAND2* at the maternal-fetal interface of eutherian mammals compared to other amniotes, with the evolution of implantation and length of embryonic retention (Marinić et al., 2021 peer-reviewed preprint). Differential expression of *HAND2* in *Chalcides ocellatus* during gestation contributed to enrichment of the GO term for epithelial tube morphogenesis (Brandley et al., 2012). This GO term was associated with eggshell gland formation in a comparative transcriptomics study on oviparous and viviparous *Phrynocephalus* lizards (Gao et al., 2019). Evidence for EDS of *HAND2* is most significant in the all-branch test, where no foreground was specified (LRT for EDS p-value= 5.90E-06; sites with ER ≥10 for positive selection= 8). However, EDS was also distinguishable from the background in several foreground tests. Ranked from most significant to least significant, EDS of *HAND2* was identified in the viviparous viperids test (LRT for EDS p-value= 1.70E-04; sites with ER ≥10 for positive selection=6), the all viviparous test (LRT for EDS p-value= 6.49E-04; sites with ER ≥10 for positive selection=6), all oviparous test (LRT for EDS p-value= 9.40E-03; sites with ER ≥10 for positive selection= 10), and oviparous viperid test (LRT for EDS p-value= 4.67E-02; sites with ER ≥10 for positive selection= 7). The only test that did not have significant EDS for *HAND2* was the viviparous natricids and elapids (LRT for EDS p-value= 0.5). Further work is necessary to determine the role of *HAND2* in squamate reproduction.

Several candidate hub genes for placenta formation in *Phrynocephalus vlangalii* (Gao et al., 2019) have evidence for EDS—*HOOK3* and *LRP6* (multiple viviparous and oviparous branch tests), *CELSR1* (all viviparous test), *ZNF462* (viviparous natricids and elapids), and *KRT80* (all oviparous). A gene with shared viviparity-specific amino acid replacements across four viviparous lizards that is expressed during gestation and involved with immune regulation (Gao et al., 2019), *NKTR*, has evidence of EDS in several tests. Subsets of genes previously associated with transitions to viviparity (Cornetti et al., 2018) also have EDS in all tests.

A previous GWAS study on a reproductively bimodal skink, *Zootoca vivipara*, associated several genes with embryonic retention and eggshell formation (Recknagel et al., 2021). Of the 288 candidates for embryonic retention included here, 107 have evidence for EDS in the all-branch test. Similarly, of the thirty-seven included candidates for eggshell formation from *Z. vivipara* (Recknagel et al., 2021), EDS was found across tests including non-overlapping sets in the oviparous tests—oviparous viperids (*LYPLA1* and *SPRED2*) and all oviparous (*ARHGAP11A, GOLGA5, IL2RB*, and *PDIA2*). These results, in conjunction with the literature (Recknagel et al., 2021), may allow researchers to explore further how squamate eggshell formation differs from birds. This will compliment recent advances looking at the presence and absence of avian-eggshell specific genes in squamate genomes (Xie et al., 2022).

Of the thirty-two differentially expressed genes during gestation in four or more squamates (Recknagel et al., 2021), eleven have EDS in the all-branch test (*ACVR2B, CDH5, EDEM3, ELL2, LMBRD2, NR4A2, RASEF, SCNN1B, SLC9A2, SMAD6*, and *TMEM181*). Of these, only *EDEM3* is significant in the all viviparous test. *EDEM3* is highly expressed during the stage of placenta formation in *Phrynocephalus vlangalii* (Gao et al., 2019); upregulated in the uterus of the chorioallantoic placenta in *Pseudemoia entrecasteauxii* (Griffith et al., 2016); and upregulated in the uterus during gestation in *Chalcides ocellatus* (Brandley et al., 2012). Interestingly, the LRT for EDS of *EDEM3* in viviparous viperids foreground branches was more significant (p-value= 1.70E-04) than in the all branch test (p-value=2.08E-03), suggesting that the all branch test may have picked up on EDS in foreground viviparous viperids. Together, this makes *EDEM3* a candidate for the evolution of the placenta, apposition of maternal and fetal tissues (Mossman 1937), in snakes—especially viviparous viperids.

Overall, this study highlights the putative role of EDS as an evolutionary driver of both oviparity and viviparity in snakes. If readers are interested in exploring the results of a specific gene, utilize HyPhy Vision (http://vision.hyphy.org). Examine genes of interest in your datasets. Echoing others (Whittington et al., 2022), I encourage future research to characterize uterine gene expression profiles in reproductively bimodal snakes during reproduction (Whittington et al., 2022). *Echis carinatus, Helicops angulatus, Protobothrops jerdonii*, and *Psammophylax variabilis* present good opportunities for this. Future studies on reproductively bimodal snakes will 1) inform how to interpret the results of this study and 2) impact how science understands parity mode evolution in Serpentes. Further research should also determine the frequency of genes under selection in snake genomes. This will enable researchers to determine if the number of genes under diversifying selection to support parity mode evolution significantly differ from what is expected by chance.

## Data Availability

Supplementary Material is available at Figshare. S1 is a folder with all gene alignments. S2 is a folder with results from BUSTED tests for episodic diversifying selection including jsons, raw text reports, and lists of genes significant for each test. S3 contains the gene trees. S4 is a folder with the gene trees. S5a is a file describing the candidate genes. S5b has the references pertinent to S5a. S6 is a file with lists of genes that are significant in multiple tests. The reads for all genomes are available on NCBI Sequence Read Archive. The genomes are available at NCBI Genome Archive.

## Acknowledgements

Special thanks to Peter Andolfatto, Cheryl Hayashi, Chris Raxworthy, and Brian Smith for their feedback on this project. Thank you to Frank Burbrink and Sean Harrington for sequencing the *Agkistrodon contortrix* genome. Thank you to the Richard Gilder Graduate School, the Department of Herpetology at the American Museum of Natural History, and the Sydney Anderson Travel Grant for funding. For loaning tissue samples, thank you to the Texas Natural History Collection at the Texas Memorial Museum, the Department of Herpetology at the California Academy of Sciences, the Division of Herpetology at the University of Kansas, the Dallas Zoo, and the Department of Herpetology at the American Museum of Natural History, stored in the Ambrose Monell Cryo Collection. Thank you as well to the Sackler Institute of Comparative Genomics for laboratory access and the New York Genome Center for genome sequencing. Thank you as well to Rachel Carroll for providing feedback on final drafts of figures.

